# *Wolbachia* infections in *Aedes aegypti* differ markedly in their response to cyclical heat stress

**DOI:** 10.1101/073106

**Authors:** Perran A. Ross, Itsanun Wiwatanaratanabutr, Jason K. Axford, Vanessa L. White, Nancy M. Endersby-Harshman, Ary A. Hoffmann

## Abstract

*Aedes aegypti* mosquitoes infected with *Wolbachia* bacteria are currently being released for arbovirus suppression around the world. Their potential to invade populations and persist will depend on interactions with environmental conditions, particularly as larvae are often exposed to fluctuating and extreme temperatures in the field. We reared *Ae. aegypti* larvae infected with different types of *Wolbachia* (*w*Mel, *w*AlbB and *w*MelPop) under diurnal cyclical temperatures. Rearing *w*Mel and *w*MelPop-infected larvae at 26-37°C reduced the expression of cytoplasmic incompatibility, a reproductive manipulation induced by *Wolbachia*. We also observed a sharp reduction in the density of *Wolbachia* in adults. Furthermore, exposure to 26-37°C over two generations eliminated both the *w*Mel and *w*MelPop infections. In contrast, the *w*AlbB infection was maintained at a high density, exhibited complete cytoplasmic incompatibility, and was transmitted from mother to offspring with a high fidelity under this temperature cycle. These findings have implications for the success of *Wolbachia* interventions across different environments and highlight the importance of temperature control in rearing.

## Introduction

*Aedes aegypti* mosquitoes transmit some of the most important arboviral diseases worldwide. They are widespread in tropical and subtropical regions (1), inhabiting urban environments where they have adapted to breed in artificial containers (2). Dengue and Zika are among the viruses they transmit and these are rapidly increasing their burden on global health. Dengue alone infects as many as 390 million people each year, and up to half of the world’s population is at risk of infection (1). Zika is an emerging threat that is experiencing an epidemic following an outbreak in Brazil in 2015 (3, 4). Efforts to reduce the spread of dengue and Zika rely on direct control of *Ae. aegypti* populations because there are no commercially available vaccines (5). Though permanent mosquito eradication is unlikely to be achieved, several genetic and biological approaches are being utilized to reduce the burden of arboviruses (6).

One such approach involves the release of *Aedes aegypti* infected with the bacterium *Wolbachia* into wild populations of mosquitoes in an effort to combat dengue and Zika (7, 8). *Wolbachia* are transmitted maternally and often manipulate the reproduction of their hosts to enhance their own transmission (9). These bacteria are of particular interest in the control of arboviral diseases as they are known to inhibit the replication of RNA viruses in insects (10). Infections of *Wolbachia* from *Drosophila melanogaster* and *Ae. albopictus* were recently introduced experimentally into *Ae. aegypti* and were found to suppress the transmission of dengue (11, 12), Zika (13, 14), chikungunya (11, 15), yellow fever (15) and West Nile viruses (16). This innate viral suppression makes *Wolbachia* a desirable alternative for arboviral control as it removes the need for mosquito eradication.

More than four *Wolbachia* infections have now been established in *Ae. aegypti* from interspecific transfers, including the *w*MelPop (17) and *w*Mel (18) infections from *D. melanogaster*, the *w*AlbB infection from *Ae. albopictus* (19), and a *w*Mel/*w*AlbB superinfection (20). These *Wolbachia* infections induce cytoplasmic incompatibility in *Ae. aegypti*, a phenomenon that results in sterility when an infected male mates with an uninfected female. *Wolbachia-*infected females therefore possess a reproductive advantage because they can produce viable offspring with both infected and uninfected males as mates (21). These infections vary considerably in their effects on the mosquito host, from the minor deleterious fitness effects of *w*Mel (22–24) to the severe longevity and fertility costs of *w*MelPop (25–27). Variability also exists in the extent to which they suppress arboviruses; infections that reach a higher density in the host tend to block viruses more effectively (12, 18, 20).

With its lack of severe fitness effects and its ability to cause cytoplasmic incompatibility, the *w*Mel infection is suitable for invading naïve mosquito populations (18). This infection has become established in multiple wild populations of mosquitoes in Queensland, Australia (7), and persists in these populations years after the associated releases ceased (22). *w*Mel is currently the favored infection for *Wolbachia* interventions on an international scale and is undergoing field release trials in Brazil, Indonesia, Vietnam and Colombia (8, 28). Cage and field trials of the *w*MelPop infection demonstrate its difficulty in invading and persisting (18, 29), though the infection could have utility in population suppression programs (30, 31) due to its detrimental effect on quiescent egg viability (25, 26). The *w*AlbB infection is yet to be released in the field but it has successfully invaded caged populations in the laboratory (19, 23).

Since *Wolbachia* were introduced into *Ae. aegypti*, the four described infections have each displayed complete cytoplasmic incompatibility and maternal inheritance in the laboratory (17–20). A high fidelity of these traits is necessary for the success of *Wolbachia* as a biological control; maternal transmission leakage and partial cytoplasmic incompatibility will increase the proportion of infected mosquitoes needed for the infection to spread, reduce the speed of invasion and prevent the infection from reaching fixation in a population (32). Some natural *Wolbachia* infections in *Drosophila* exhibit perfect maternal inheritance and complete cytoplasmic incompatibility in the laboratory, but display incomplete fidelity under field conditions (33, 34).

The effects of *Wolbachia* on reproduction can depend on the density of *Wolbachia* in mosquito cells. In insects other than *Ae. aegypti*, a decline in *Wolbachia* density can reduce the degree of male-killing (35), feminization (36), parthenogenesis (37) and cytoplasmic incompatibility (38, 39). Incomplete cytoplasmic incompatibility occurs when some sperm cysts in the testes are not infected with *Wolbachia* (40, 41). To ensure the transmission of *Wolbachia* to all offspring, densities must exceed a threshold in the ovaries (42). Viral protection by *Wolbachia* is also density dependent, with higher densities in the host generally resulting in greater protection (43, 44). However, environmental conditions such as temperature (45, 46), nutrition (47, 48) and pathogen infection (49, 50) are known to modulate *Wolbachia* densities in other insects. Given the importance of bacterial density in determining *Wolbachia*’s reproductive effects (cytoplasmic incompatibility and maternal transmission fidelity), fitness costs and viral blocking effects, work is needed to determine if environmental effects play a role in modulating densities in experimental infections of *Ae. aegypti*.

*Ae. aegypti* larvae often experience large diurnal fluctuations of temperature in nature, particularly in small containers of water and in habitats exposed to direct sunlight (51, 52). While the thermal limits of *Ae. aegypti* are generally well understood (53–55), research has not assessed *Wolbachia*’s reproductive effects in *Ae. aegypti* at the high temperatures they can experience in the field. Ulrich and others (56) recently demonstrated that the density of *w*Mel in *Ae. aegypti* decreased sharply when larvae experienced diurnally cycling temperatures of 28.5°C to 37.5°C during development. This suggests that the reproductive effects of *Wolbachia* could also be altered if infected larvae develop under similar conditions in the field.

We explored the hypothesis that the reproductive effects of *Wolbachia* infections could be diminished if *Ae. aegypti* experience stressful, high thermal maxima within a large diurnal cyclical temperature regime during development. We tested three *Wolbachia* infections: *w*Mel, *w*MelPop and *w*AlbB, for their maternal transmission fidelity and ability to cause cytoplasmic incompatibility. We show for the first time that cyclical temperatures reaching a maximum of 37°C reduce the expression of cytoplasmic incompatibility in the *w*Mel and *w*MelPop infections of *Ae. aegypti*. We also find a greatly diminished *Wolbachia* density under these conditions. Exposing *w*Mel and *w*MelPop-infected mosquitoes to this regime over two generations cures *Wolbachia* entirely. Conversely, the *w*AlbB infection is more stable in terms of its reproductive effects and density under cyclical temperatures. These findings suggest the need for multiple infection types suitable for different conditions when using *Wolbachia* infections in biological control strategies.

## Results

### Maximum daily temperatures of 37°C during development reduce the hatch rate of *w*Mel-infected eggs

We compared the hatch rate of eggs from crosses between *Wolbachia*-infected *Ae. aegypti* females and *Wolbachia*-infected males reared under cyclical temperatures. Larvae of both sexes were reared in incubators set to cycle diurnally between a minimum of 26°C and a maximum of either 26°C, 32°C, 34.5°C or 37°C (Figure S1), and crosses were then conducted at 26°C. We observed a sharp decrease in the hatch rate of eggs when *w*Mel-infected mosquitoes were reared at 26-37°C compared to 26°C (Mann-Whitney U: Z = 2.802, P =0.005), but found no effect of rearing temperature on hatch rate for the *w*AlbB (Kruskal-Wallis *χ*^2^ =2.587, df =3, P =0.460) or *w*MelPop (*χ*^2^ =1.687, df =3, P =0.640) infections (Figure 1.). We hypothesized that reduced hatch rate in *w*Mel-infected mosquitoes could reflect the loss of *Wolbachia* infection under heat stress, leading to partial cytoplasmic incompatibility.

**Figure 1.**
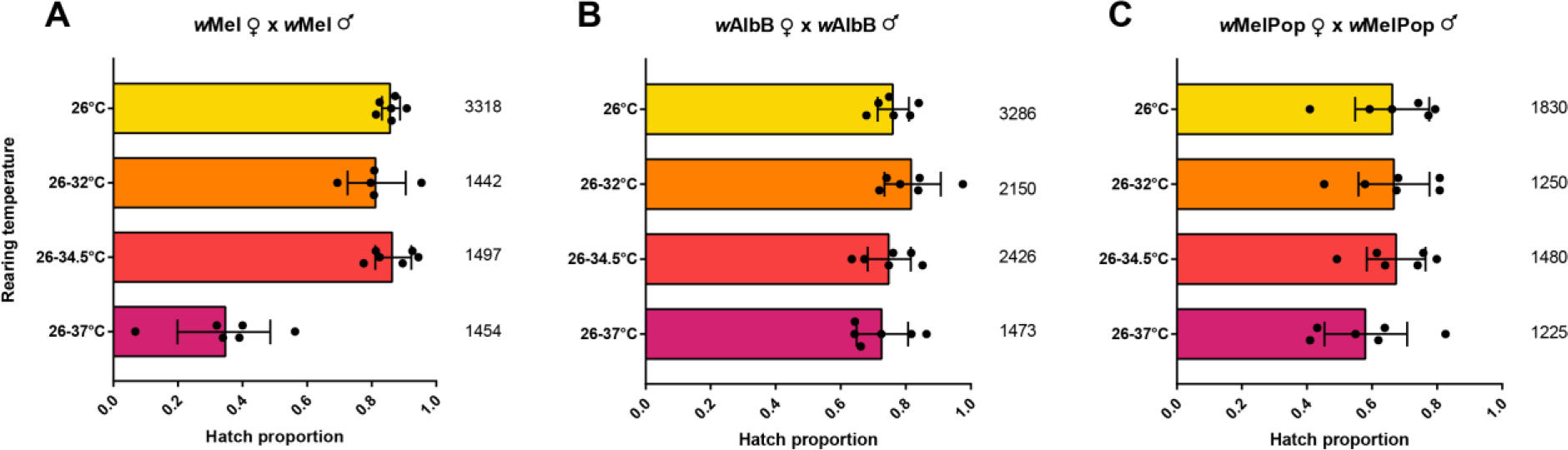
Proportion of eggs hatched from *Wolbachia*-infected females crossed to *Wolbachia*-infected males reared at different cyclical temperatures for the (A) *w*Mel, (B) *w*AlbB and (C) *w*MelPop infections. Both sexes were reared under the same temperature regime and then crossed together at 26°C. Each data point shows the proportion of eggs hatched from a cage of 7 females and 7 males. Numbers for each bar denote the total number of eggs scored per cross. Error bars show 95% confidence intervals.

### *Wolbachia* density is reduced in *w*Mel and *w*MelPop, but not *w*AlbB-infected adults reared under cyclical temperatures of 26-37°C

We wanted to see if a reduction in *Wolbachia* density could explain the reduced hatch rate of *w*Mel-infected eggs. We measured the density of *Wolbachia* in adults infected with *w*Mel, *w*AlbB and *w*MelPop when reared at either 26°C, 26-32°C or 26-37°C. The density of *w*Mel did not differ significantly between 26°C and 26-32°C for either males (Mann-Whitney U: Z =1.190, P =0.234) or females (Z =1.112, P =0.267), but sharply decreased at 26-37°C in both sexes (Figure 2). The density in females reared at 26°C (mean ± SD =3.56 ± 1.87, n =29) was 14.75-fold higher than those reared at 26-37°C (0.24 ± 1.04, n =30, Z =6.239, P < 0.0001). For males the difference between 26°C (4.65 ± 2.71, n =29) and 26-37°C (0.027 ± 0.025, n =30) was 174.73-fold (Z =6.688, P < 0.0001). For *w*MelPop, female *Wolbachia* density at 26°C (mean ± SD =84.60 ± 89.19, n =30) was 268.34-fold higher than those reared at 26-37°C (0.32 ± 0.66, n =30, Z =6.631, P < 0.0001)., while males reared at 26°C (45.62 ± 32.25, n =30) had a 73.37-fold higher density than males reared at 26-37°C (0.62 ± 1.76, n =30, Z =6.542, P < 0.0001). In contrast, there was no significant difference in *w*AlbB density between 26°C and 26-37°C for both females (Z =0.47 P =0.638) and males (Z =1.678, P =0.093). However, there was a significant effect of temperature overall due to an increased density at 32°C in both females (Kruskal-Wallis *χ*^2^ =7.826, df =2, P =0.020) and males (*χ*^2^ =16.311, df =2, P =0.0003).

**Figure 2.**
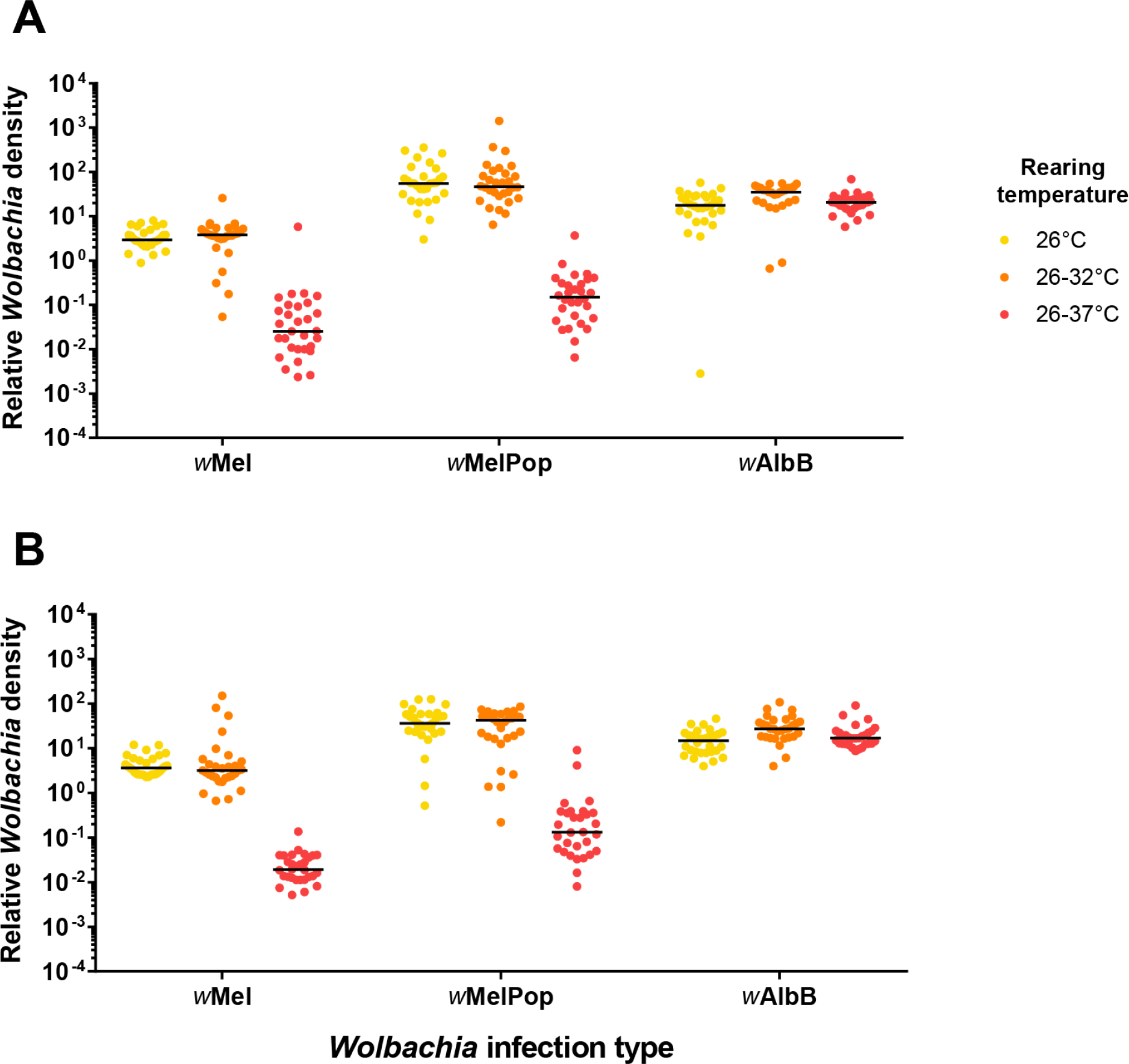
Relative *Wolbachia* density in (A) female and (B) male adults reared at a constant 26°C, cycling 26-32°C or cycling 26-37°C. Each mosquito was tested with mosquito-specific and *Wolbachia*-specific markers to obtain crossing point values (see “ *Wolbachia* quantification”). Differences in crossing point between the two markers were transformed by 2^n^ to obtain relative *Wolbachia* densities. Each data point represents the average of three technical replicates.

### Cytoplasmic incompatibility is partially lost in *w*Mel and *w*MelPop, but not *w*AlbB-infected adults reared under cyclical temperatures of 26-37°C

Crosses between uninfected female and *Wolbachia*-infected male *Ae. aegypti* produce no viable offspring under standard laboratory conditions due to cytoplasmic incompatibility (17–19). We hypothesized that reduced *Wolbachia* densities in infected males reared at 26-37°C would coincide with reduced fidelity of cytoplasmic incompatibility. Incomplete cytoplasmic incompatibility leads to some viable progeny when infected males mate with uninfected females (33). We crossed *w*Mel, *w*AlbB and *w*MelPop males reared at 26°C and 26-37°C to uninfected females reared at 26°C, and scored the proportion of eggs that hatched (Figure 3A). 245 larvae hatched from 1747 eggs (14.02%) across all replicates when *w*Mel-infected males were reared at 26-37°C. Conversely, we observed complete sterility when males were reared at 26°C (Mann-Whitney U: Z =2.802, P =0.005). We also observed incomplete cytoplasmic incompatibility in the *w*MelPop infection; 301 larvae hatched from 1846 eggs (16.31%) when males were reared at 26-37°C, but no larvae hatched when males were reared at 26°C (Z =2.802, P =0.005). In contrast to *w*Mel and *w*MelPop, no eggs hatched from uninfected females that were mated to *w*AlbB-infected males reared under either regime (Z =0.080, P =0.936). The cytoplasmic incompatibility induced by *w*AlbB therefore appears to be stable under these conditions.

We also scored the hatch rate of *Wolbachia*-infected females reared under a cycling 26-37°C when crossed to infected males reared at 26°C (Figure 3B). We hypothesized that reduced *Wolbachia* densities in the female could restore cytoplasmic incompatibility in this cross. For the *w*Mel infection, mean hatch rates were drastically reduced to 22.7% in infected females reared at 26-37°C compared to 85.7% when reared at 26°C (Mann-Whitney U: Z =2.802, P =0.005). Conversely, we found no effect on the *w*MelPop (Z =0.400, P =0.689) and *w*AlbB (Z =0.560, P =0.575) infections; females possessed similar hatch rates regardless of the rearing temperature. Taken together, these results show that a cyclical rearing regime reaching a maximum of 37°C reduces both the ability of *w*Mel-infected males to induce cytoplasmic incompatibility and the ability of *w*Mel-infected females to retain compatibility.

**Figure 3.**
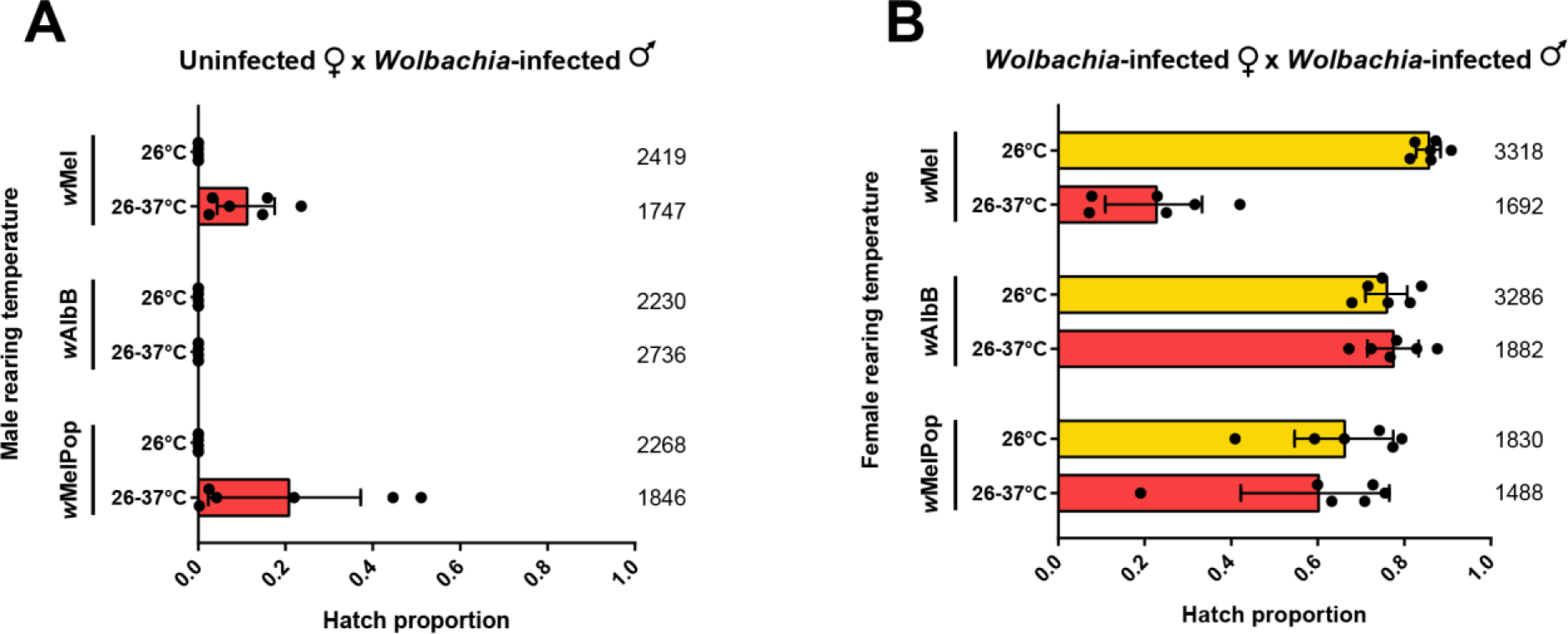
(A) Proportion of eggs hatched from uninfected females reared at 26°C and *Wolbachia-*infected males reared at either 26°C or a cycling 26-37°C. (B) Proportion of eggs hatched from *Wolbachia*-infected females reared at either 26°C or 26-37°C and *Wolbachia*-infected males of the same infection type reared at 26°C. For both sets of crosses, adults were mated at 26°C after a period of maturation. Each data point shows the proportion of eggs hatched from a cage of 7 females and 7 males. Numbers for each bar denote the total number of eggs scored per cross. Error bars show 95% confidence intervals.

### The *w*Mel and *w*MelPop infections are lost, and *w*AlbB exhibits incomplete maternal transmission fidelity at 26-37°C

We tested the ability of *w*Mel, *w*AlbB and *w*MelPop-infected females to transmit *Wolbachia* to their offspring when their entire lifecycle occurred at either a constant 26°C or a cycling 26-37°C. Females from each infection type were crossed to uninfected males, and their progeny were reared to the 4^th^ instar at the same temperature as the mother. *w*Mel and *w*AlbB-infected females transmitted the infection to all of their offspring at 26°C. The *w*MelPop infection was also transmitted with a high fidelity at 26°C, though a single *w*MelPop-infected female produced two uninfected progeny (Table 1). In contrast, the *w*Mel and *w*MelPop infections were lost completely when mothers and offspring were maintained at 26-37°C; all progeny were conclusively uninfected with *Wolbachia*. The *w*AlbB infection was transmitted to the majority of offspring at 26-37°C, but 11.5% lost the infection (Table 1).

**Table 1.** Proportion of *Wolbachia*-infected offspring produced by *w*Mel, *w*MelPop and *w*AlbB-infected mothers when maintained at a constant 26°C or a cycling 26-37°C. Ten progeny from eight mothers, for a total of 80 progeny, were tested per treatment.

| *Wolbachia*<br>infection type | Temperature | Maternal<br>transmission rate | Binomial confidence interval<br>(lower 95%, upper 95%) |
| --- | --- | --- | --- |
| *wMel* | 26°C | 1 | 0.955, 1 |
|  | 26-37°C | 0 | 0, 0.045 |
| *wMelPop* | 26°C | 0.975 | 0.912, 0.997 |
|  | 26-37°C | 0 | 0, 0.045 |
| *wAlbB* | 26°C | 1 | 0.955, 1 |
|  | 26-37°C | 0.885 | 0.792, 0.946 |

## Discussion

We demonstrate for the first time that the *w*Mel and *w*MelPop infections of *Ae. aegypti* exhibit reduced cytoplasmic incompatibility when immature stages experience cyclical temperatures of 26-37°C during development, in contrast to the *w*AlbB infection. We also show that these infections are lost completely when infected mosquitoes experience these conditions over their entire lifecycle. *w*Mel infected mosquitoes are currently being deployed in several countries for the control of arboviruses (28). Immature *Ae. aegypti* may experience extreme temperatures in the field (31, 52), and the thermal sensitivity of the *w*Mel and *w*MelPop infections could therefore reduce their ability to establish and persist in natural populations. The *w*AlbB infection retains its ability to induce complete cytoplasmic incompatibility under the same conditions, while maternal transmission fidelity remains relatively high. Densities of *w*AlbB are also stable, suggesting that it will also provide effective arboviral protection (20, 57). The robustness of *w*AlbB when exposed to high maximum temperatures could make this infection more suited for field release in environments where temperatures in breeding sites fluctuate in comparison to *w*Mel.

High temperatures have been known for some time to have a negative effect on *Wolbachia*. In other arthropods, high temperatures can reduce the density of *Wolbachia* in its host (45, 58–60), weaken the reproductive effects induced by *Wolbachia* (38, 61–65) and even eliminate *Wolbachia* entirely (61, 62, 66–68). Only recently have the effects of temperature been characterised in experimental *Wolbachia* infections of *Ae. aegypti*. Ye and others (69) reared *w*Mel-infected larvae under diurnally cycling temperatures and assessed their vector competence and *Wolbachia* density. They concluded that the *w*Mel infection should remain robust in terms of its ability to reduce dengue transmission under field conditions in Cairns, Australia. However, the authors only tested temperatures reaching a maximum of 32°C; we observed no effect on *Wolbachia* density or hatch rate under similar conditions. In nature, larvae and pupae are restricted to aquatic environments where average maximum temperatures can reach 37°C during the wet season in Cairns (52). Here we employed a larger temperature range to better reflect natural conditions in the field. Although we did not test vectorial capacity directly, we observed a greatly reduced density of *w*Mel in adults when larvae experienced a maximum temperature of 37°C. These conditions will likely affect the viral suppression induced by *w*Mel as the ability of *Wolbachia* to interfere with transmission relies on high densities in relevant tissues (43, 70).

In the majority of our experiments we exposed larvae to cyclical temperatures while maintaining adults and eggs at 26°C. However, we observed that when all life stages were maintained at 26-37°C for almost two generations, the *w*Mel and *w*MelPop infections were eliminated. The *w*AlbB infection also exhibited some maternal transmission leakage despite maintaining high densities and complete cytoplasmic incompatibility when only larvae were exposed. This suggests that both the duration of exposure and the maximum temperature reached will affect *Wolbachia* density. Ulrich and others (56) provide additional evidence that the timing of heat stress is important; lowest *w*Mel densities corresponded with the longest stress duration in immature *Ae. aegypti*, and densities varied considerably depending on their developmental stage at the time of exposure. More work is needed in these areas particularly as conditions and responses in the field are likely to be diverse.

We find that the *w*Mel and *w*MelPop infections differ markedly from *w*AlbB in their response to heat stress; to our knowledge this is the first comparison of high temperature responses between multiple *Wolbachia* infections. Differences in heat tolerance could result from different evolutionary histories; *w*Mel and *w*MelPop are nearly genetically identical (71, 72) and originate from the same host, *D. melanogaster* (73, 74). *w*AlbB occurs naturally in *Ae. albopictus*, a mosquito native to south-east Asia (75, 76); this infection may have evolved a relatively higher heat tolerance in response to the temperatures experienced by *Ae. albopictus* in its historical distribution. *w*AlbB density decreases only slightly when naturally infected *Ae. albopictus* are reared at a constant 37°C (45, 77). The effects of high temperatures on the density of *w*Mel and *w*MelPop in their natural host are however unknown.

Whether there is an influence of the host on *Wolbachia*’s thermal tolerance requires further investigation.

The differential responses of *Wolbachia* infection types under heat stress may arise from factors other than their ability to tolerate high temperatures. *Wolbachia* densities can be influenced by interactions with WO, temperate bacteriophage which infect *Wolbachia* (78). Temperate phage undergo lysogenic and lytic cycles, the latter of which can be induced by heat shock (59, 79) During the lytic cycle, phage replicate and infect new *Wolbachia* cells, potentially reducing densities of *Wolbachia* through cell lysis (80). High densities of lytic phage reduce the density of *Wolbachia* and the strength of cytoplasmic incompatibility in the parasitoid wasp *Nasonia vitripennis* (81). Therefore, high temperatures may reduce *Wolbachia* densities in *Ae. aegypti* through the same mechanism. WO phage infect *w*Mel (82) and *w*AlbB (50, 83) in their native hosts, but it is unknown if they persist following transfer to *Ae. aegypti*, though WO phage can be maintained upon interspecific transfer of *Wolbachia* in moths (84). This too requires further investigation.

While the mechanism for the loss of *Wolbachia* at high temperatures is unknown, our results strongly suggest that the ability of *w*Mel and *w*MelPop-infected *Ae. aegypti* to invade and persist in natural populations will be adversely affected by heat. We observed reduced cytoplasmic incompatibility and maternal transmission fidelity at cyclical temperatures approximating breeding containers in the field; constant high temperatures are therefore not needed to have adverse effects on *Wolbachia*. Incomplete cytoplasmic incompatibility and/or maternal transmission fidelity of *Wolbachia* will reduce the speed of invasion, increase the minimum infection threshold required for invasion to take place and decrease the maximum frequency that can be reached in a population (32). Maximum daily temperatures of larval mosquito habitats in nature can reach or exceed the maximum temperature tested in this study (52, 85, 86) and this should be a careful consideration for additional research in this area. Though *Wolbachia* densities may partially recover if adults can avoid extreme temperatures (56), the loss of cytoplasmic incompatibility can still occur even when adults are returned to low temperatures for several days before mating, as we demonstrate here. These findings could help explain the lack of invasiveness by the *w*Mel infection in some tropical locations where upper extremes are common (28). Mosquito suppression strategies which use *Wolbachia*-infected males as a sterile insect may also be impacted by temperature but this work suggests males reared in the laboratory at lower temperatures are more likely to succeed.

As releases of *Ae. aegypti* infected with *w*Mel are currently underway in several countries, researchers should assess the impact of heat stress on *Wolbachia* infections in the field. Our findings emphasize the need for further characterization of current *Wolbachia* infections under a range of temperature conditions, particularly in terms of the duration of exposure to extreme temperatures and the effects across generations An enormous diversity of *Wolbachia* strains exist in nature (87); alternative strains, or current infections selected for increased thermal tolerance, should be considered. Our results also highlight the importance of temperature control in the laboratory rearing of *Wolbachia*-infected insects. Heat stress could be used to cure the *w*Mel and *w*MelPop infections from mosquitoes in order to study their effects (68) as an alternative to tetracycline (88). A better understanding of the response of *Wolbachia* infections to varying environmental conditions is required particularly in the context of laboratory rearing and in their application an arboviral biocontrol agent in the field.

## Methods

### Colony maintenance and *Wolbachia* infections

Uninfected *Aedes aegypti* mosquitoes were collected from Townsville, Queensland, in November 2015 and maintained in a temperature controlled insectary at 26°C ± 1°C according to methods described by Axford and others (23). *Aedes aegypti* with the *w*Mel, *w*MelPop and *w*AlbB infections of *Wolbachia* were derived from lines transinfected previously (17–19). Females from all *Wolbachia-* infected lines were crossed to males from the Townsville line for three generations in succession to control for genetic background. Female mosquitoes were blood fed on the forearms of human volunteers. Blood feeding on human subjects was approved by the University of Melbourne Human Ethics Committee (approval 0723847). All volunteers provided informed written consent.

### Rearing at cyclical temperatures

Larvae for all experiments were reared in incubators (PG50 Plant Growth Chambers, Labec Laboratory Equipment, Marrickville, NSW, Australia) set to a constant 26°C or to one of the following cyclical temperatures: 26-32°C, 26-34.5°C and 26-37°C at a 12:12 light: dark photoperiod. Cycling incubators were set to maintain 26°C during the dark period and the maximum temperature during light, with 12 hours at each temperature. Water temperatures were monitored by placing data loggers (Thermochron; 1-Wire, iButton.com, Dallas Semiconductors, Sunnyvale, CA, USA) in sealed glass vials, which were submerged in plastic trays (11.5 × 16.5 × 5.5 cm) filled with 500 mL of water identical to larval rearing trays. Temperature was measured at 30 minute intervals. Representative daily temperature fluctuations that occurred in each incubator for the duration of the experiments are shown in Figure S1. Rearing at cyclical temperatures of 26-32°C or 26-37°C decreased the wing length of adults (Figure S2), suggesting they were heat stressed.

For each experiment, eggs from the uninfected, *w*Mel, *w*MelPop and *w*AlbB lines were hatched synchronously in 3 L trays of RO water at 26°C. Hatching trays were transferred to incubators within two hours of hatching. Larvae were provided TetraMin® tropical fish food tablets (Tetra, Melle, Germany) *ad libitum* and maintained at a controlled density of 100 larvae in 500 mL water. Temperatures in each incubator deviated by up ±0.5°C from the set-point, depending on the location of data loggers. We randomised the position of rearing trays within incubators and frequently moved them to different positions to account for positional effects.

### Hatch rate and cytoplasmic incompatibility

Crosses between *Wolbachia* infection types were conducted to determine the proportion of viable offspring from parents reared at different cyclical temperatures. Pupae were sexed according to size (females are larger than males) and added to 12 L cages held at 26°C ± 1°C within 24 hours of eclosion after confirming their sex. Sexes, infection types and adults reared at each temperature were maintained in separate cages. Adults were allowed to mature and acclimatise to 26°C for at least 48 hours; crosses were conducted only when all adults were at least 48 hours old as development times varied between sexes and rearing temperatures. After the period of maturation, 7 males and 7 females from their respective infection type were aspirated into 1.5 L cages and allowed to mate for 3 days. Each cross was comprised of 6 replicate cages; the combinations of sex, rearing temperature and *Wolbachia* infection status for each cross are described in the results section. Each cage was provided with water for the duration of the experiment, and sugar until 24 h prior to blood feeding. Females were provided a blood meal through mesh on the side of each cage until all females had fed to repletion. Multiple human volunteers were used, with one volunteer per replicate cage. Pill cups were filled with 25 mL of water and lined with filter paper (Whatman® 90mm qualitative circles, GE Healthcare Australia Pty. Ltd., Parramatta, New South Wales, Australia) and provided as an oviposition substrate. Eggs laid on filter papers were collected daily, dried on paper towel and photographed with a digital camera. The number of eggs laid was determined with a clicker counter. Eggs were hatched in containers of 200 mL of water four days after collection, and larvae were reared to the 3^rd^ instar. Hatch proportions were defined as the number of larvae counted, including larvae that hatched precociously (visible on the filter papers).

### *Wolbachia* quantification

The density of *Wolbachia* in adults reared at cyclical temperatures was determined for the *w*Mel, *w*MelPop and *w*AlbB infections. We reared three trays of 100 larvae per infection type at 26°C, 26-32°C and 26-37°C (see “Rearing at cyclical temperatures”). Eclosing adults were collected daily at noon and stored in absolute ethanol for DNA extraction. We selected 10 males and 10 females at random per tray for *Wolbachia* quantification. DNA extraction and *Wolbachia* quantification were conducted according to methods described previously (22, 23, 89). DNA from adults with both wings removed was extracted using 150 µL of 5% Chelex® 100 resin (Bio-Rad Laboratories, Hercules, CA). We used a LightCycler 480 system (Roche Applied Science, Indianapolis, IN) to amplify *Ae. aegypti-* specific (*aRpS6*) and *Wolbachia*-specific (*w*Mel, *w*AlbB or *w*MelPop) genes. Three technical replicates of the *aRpS6* and *Wolbachia*-specific markers were completed for each mosquito; differences in crossing point between the two markers were averaged to obtain an estimate of *Wolbachia* density. These values were then transformed by 2^n^ to obtain relative *Wolbachia* densities.

### Maternal transmission of *Wolbachia*

We tested the ability of *w*Mel, *w*MelPop and *w*AlbB-infected females to transmit *Wolbachia* infections to their offspring. *Wolbachia*-infected females were reared from the egg stage in incubators set to a constant 26°C or a cycling 26-37°C (see “Rearing at cyclical temperatures”) and crossed to uninfected males. Females were blood-fed *en masse* and isolated in 70mL plastic cups filled with 20mL of water and lined with a 2 × 12 cm strip of sandpaper (Norton® Master Painters P80 sandpaper, Saint-Gobain Abrasives Pty. Ltd., Thomastown, Victoria, Australia). Eggs from each female were hatched by adding an additional 10 mL of water to the plastic cup in order to submerge the eggs. TetraMin® was provided *ad libitum*. Progeny were reared to 3^rd^ or 4^th^ instar, stored in ethanol, then tested for the presence and density of *Wolbachia*(see “ *Wolbachia* quantification”). We scored 10 offspring from 8 females per infection type at each temperature. Note that mothers and offspring were maintained in their respective incubators (26°C or 26-37°C) for the entire duration of the experiment, including egg and adult stages.

### Statistical analyses

All analyses were conducted using SPSS statistics version 21.0 for Windows (SPSS Inc, Chicago, IL). Hatch proportions and *Wolbachia* densities were not normally distributed according to Shapiro-Wilk tests, therefore we analyzed all data with nonparametric Kruskal-Wallis and Mann-Whitney U tests.

## Acknowledgements

We thank Elizabeth Valerie, Shani Wong, Michael Ørsted, Ashley Callahan and Ellen Cottingham for providing technical assistance with experiments. We thank Chris Paton and Scott Ritchie for providing field-collected mosquito eggs. We also thank Peter Kriesner and Gordana Rašić for valuable discussions.

